# Nuclear hormone receptors promote gut and glia detoxifying enzyme induction and protect *C. elegans* from the mold *P. brevicompactum*

**DOI:** 10.1101/2021.07.15.452486

**Authors:** Sean W. Wallace, Malcolm C. Lizzappi, Hong Hur, Yupu Liang, Shai Shaham

## Abstract

Animals encounter microorganisms in their habitats, adapting physiology and behavior accordingly. The nematode *Caenorhabditis elegans* is found in microbe-rich environments; however, its responses to fungi are not extensively studied. Here we describe interactions of *C. elegans* and *Penicillium brevicompactum*, an ecologically-relevant mold. Transcriptome studies reveal that co-culture upregulates stress-response genes, including xenobiotic metabolizing enzymes (XMEs), in *C. elegans* intestine and AMsh glial cells. The nuclear hormone receptors (NHR) NHR-45 and NHR-156 are key induction regulators, and mutants that cannot induce XMEs in the intestine when exposed to *P. brevicompactum* experience mitochondrial stress and exhibit developmental defects. Different *C. elegans* wild isolates harbor sequence polymorphisms in *nhr-156*, resulting in phenotypic diversity in AMsh glia responses to microbe exposure. We propose that *P. brevicompactum* mitochondria-targeting mycotoxins are deactivated by intestinal detoxification, allowing tolerance to moldy environments. Our studies support the idea that *C. elegans* NHR gene expansion/diversification underlies adaptation to microbial environments.

## INTRODUCTION

A number of experimental paradigms have been established to study the interactions between *C. elegans* and model microbial pathogens (Jiang and Wang, 2018; Kim and Ewbank, 2018; Kim and Flavell, 2020; Madende et al., 2020; Radeke and Herman, 2021; Troemel, 2016). Recent work has also explored the interactions between *C. elegans* and the microbial components of its natural substrates in the wild (Schulenburg and Félix, 2017; Zhang et al., 2017), with most of these studies focusing on bacteria. More than 100 mycotoxins have been identified with nematocidal activities (Li et al., 2007), highlighting the notion that to survive in its habitat, *C. elegans* also possess mechanisms to counter fungi. The *Penicillium* genus includes a large number of abundant fungal species, and the mold *Penicillium brevicompactum* is found naturally in substrates for wild isolates of *C. elegans* (Dirksen et al., 2016). Interactions between *C. elegans* and fungi have not been extensively studied, and effects of *Penicillium* species, in particular, on *C. elegans* are virtually unexplored (Huang et al., 2014; Visagie et al., 2014).

Xenobiotic metabolizing enzymes (XMEs) are key detoxification proteins that include phase I enzymes of the cytochrome P450 (*cyp*) family, and phase II enzymes of the UDP-glucoronosyltransferase (*ugt*) and glutathione-s-transferase (*gst*) families (Hartman et al., 2021). XME functions have been extensively explored in the mammalian liver, where they play a major role in detoxification. XME responses are induced by the nuclear hormone receptors CAR and PXR following drug exposure (Wallace and Redinbo, 2012). The nuclear hormone receptor family in *C. elegans* (NHR) has undergone massive expansion and sequence diversification (Taubert et al., 2011). NHR genes promote responses to microbe exposure and environmental stress (Jones et al., 2013; Mao et al., 2019; Otarigho and Aballay, 2020; Park et al., 2018; Peterson et al., 2019; Rajan et al., 2019; Shomer et al., 2019; Wani et al., 2021; Ward et al., 2014; Yuen and Ausubel, 2018). Induction of XMEs has also been observed in *C. elegans* in response to exposure to several model pathogens (Engelmann et al., 2011).

Here, we establish an assay to characterize the interactions between *C. elegans* and *P. brevicompactum*. We show that NHRs mediate *P. brevicompactum*-dependent induction of XMEs in the intestine and in glia, and demonstrate a role for NHRs in protecting *C. elegans* from the toxic effects of *P. brevicompactum* exposure.

## RESULTS

### *Penicillium brevicompactum* exposure induces expression of stress response genes including xenobiotic metabolizing enzymes

To establish an assay for investigating interactions between *C. elegans* and fungal molds, we sequenced the internal transcribed spacer (ITS) rDNA regions (Schoch et al., 2012) of moldy contaminants occurring spontaneously on NGM (nematode growth medium) plates in our laboratory. *Penicillium brevicompactum* (ATCC #9056) was identified as a common contaminant, and chosen for further study because, in addition to growing readily on NGM plates suitable for *C. elegans* co-culture, it is found naturally in substrates for wild isolates of *C. elegans* (Dirksen et al., 2016). To characterize transcriptional changes in *C. elegans* following *P. brevicompactum* exposure, whole-animal RNA was extracted and sequenced from young adult nematodes grown in the presence of the standard food source *E. coli* OP50, with or without *P. brevicompactum* co-culture. We identified 100 genes that are significantly upregulated following *P. brevicompactum* exposure (fold change > 2, p-adj < 0.05, Tables S1 and S2). Functional analysis using WormCat (Holdorf et al., 2019) showed that more than one third of these are involved in stress responses (Figure 1A). In particular, statistical enrichment analysis revealed that genes belonging to the stress response sub-categories ‘detoxification’, ‘pathogen’, and ‘heavy metal’, are significantly enriched in our data set (Figure 1B). As detoxification genes are the most abundant stress-gene class modulated by *P. brevicompactum* exposure, we focused on these for further characterization.

**Figure 1.**
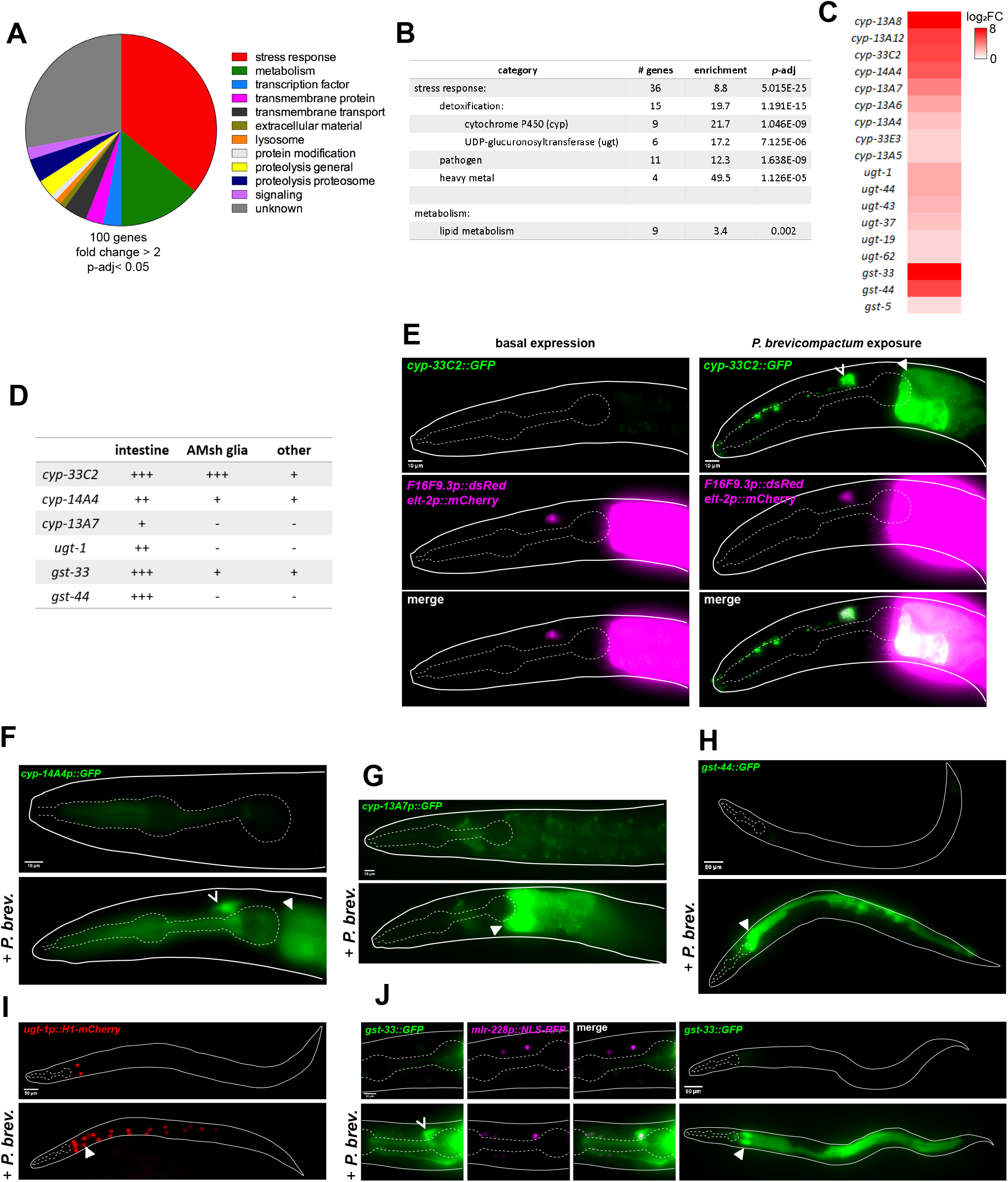
*Penicillium brevicompactum* induces expression of stress response genes including xenobiotic metabolizing enzymes (XMEs) in *C. elegans*. (A) Classification of *P. brevicompactum*-induced genes. (B) Statistically-enriched gene categories. (A) and (B) based on analysis using WormCat (Holdorf et al., 2019). (C) Heat map of XMEs induced by *P. brevicompactum*. (D) Summary of confirmed *P. brevicompactum*-induced genes. (E) Expression of *cyp-33C2::GFP* with *F16F9.3p::dsRed* (AMsh glia) and *elt-2p::mCherry* (intestine). (F-J) expression of additional XMEs listed in (D). *mir-228p::NLS-RFP* is pan-glial. Open arrowhead, AMsh glia, closed arrowhead, intestine.

Xenobiotic metabolizing enzymes (XMEs) are major detoxification proteins encoded by cytochrome P450 (*cyp*), UDP-glucoronosyltransferase (*ugt*), and glutathione-s-transferase (*gst*) gene families (Hartman et al., 2021). 18 of the *P. brevicompactum*-induced genes encode XMEs (Figure 1C). To confirm *P. brevicompactum*-dependent induction, and to identify sites of expression, we examined fluorescent transcriptional reporters for six highly-induced XMEs. All showed robust induction, validating our RNAseq data (Figure 1D-J). XME induction was consistently observed in the intestine, in line with previous reports of detoxification functions in this tissue (Hartman et al., 2021). XME induction was also observed in other tissues, and most consistently in AMsh glial cells, which are components of the major sensory organ in the head of the animal (Singhvi and Shaham, 2019). Induction of XMEs in AMsh glia is particularly interesting given that XMEs are highly expressed in sustentacular cells, the analogous cell type in the mammalian olfactory epithelium (Heydel et al., 2013).

Overall, our sequencing data demonstrates that exposure of *C. elegans* to *P. brevicompactum* causes significant changes in gene expression, strongly suggesting a biologically meaningful response to this pathogen. Furthermore, the significant enrichment of stress response genes, and specifically of detoxification genes, suggests that *P. brevicompactum* may constitute a toxic species for *C. elegans*.

### The nuclear hormone receptors *nhr-45* and *nhr-156* are required for XME induction

To investigate the molecular mechanism through which XMEs are induced by *P. brevicompactum*, we initially looked at well-characterized innate immune genes of the p38 MAP kinase pathway (Kim et al., 2002), and at cilia genes that are required for chemosensation (Inglis et al., 2007). Mutations in neither gene class affects XME induction (Figure S1A).

In light of the established role of CAR and PXR receptors in the induction of XMEs in the mammalian liver (Wallace and Redinbo, 2012), we next screened available *C. elegans nhr* RNAi clones for effects on *P. brevicompactum*-dependent induction of *cyp-33C2::GFP*, an XME reporter we developed (Figure 1E). This screen revealed that *nhr-45* is required for *cyp-33C2* induction in both intestine and AMsh glia, while *nhr-156* is partially required for *cyp-33C2* induction in AMsh glia only. These results were confirmed using a panel of loss-of-function mutants either previously identified, or generated in this study (Figure 2A-C).

**Figure 2.**
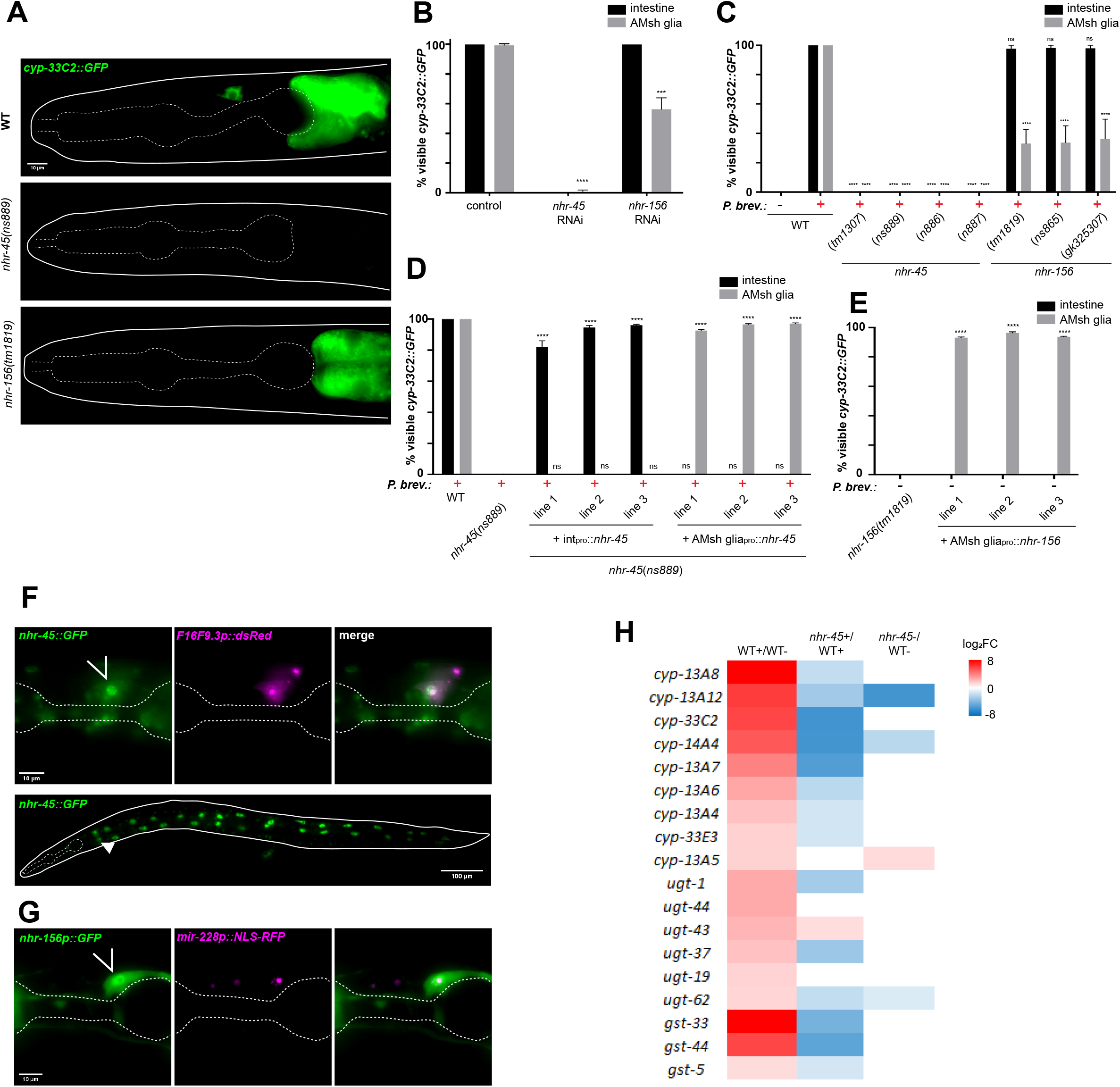
*nhr-45* and *nhr-156* regulate induction of XMEs. (A) *cyp-33C2::GFP* induction following *P. brevicompactum* exposure. (B) *cyp-33C2::GFP* induction in intestine and AMsh glia in RNAi-treated worms. **** *p* < 0.0001, *** *p* < 0.001 (t-test). Note that *p*-values can not be calculated for the intestine because there is no variation between replicates. (C) *cyp-33C2::GFP* induction in intestine and AMsh glia. **** *p* < 0.0001 (ANOVA compared to WT+*P. brev.*). (D) *cyp-33C2::GFP* induction in intestine and AMsh glia. Intpro is *elt-2p* for intestinal rescue, AMsh gliapro is *F16F9.3p* for glial rescue. **** *p* < 0.0001 (ANOVA compared to *nhr-45(ns889)*). (E) Basal expression of *cyp-33C2::GFP* in the absence of *P. brevicompactum.* **** *p* < 0.0001 (ANOVA). (F) Expression of *nhr-45::GFP* with *F16F9.3p::dsRed* (AMsh glia). Open arrow, AMsh glia, closed arrows, intestine. (G) expression of *nhr-156p::GFP* with *mir-228p::NLS-RFP* (pan-glia). Open arrow, AMsh glia. (H) Heat map of expression changes of the 18 *P. brevicompactum*-induced XMEs in *nhr-45* partial loss-of-function mutants. +, *P. brevicompactum* exposure. −, controls samples not exposed.

The two NHRs we identified function cell-autonomously to induce *cyp-33C2* expression. Introduction of an *nhr-45* cDNA under the control of intestine- or AMsh glia-specific promoters into *nhr-45* mutant animals is sufficient to rescue *cyp-33C2::GFP* induction defects in the corresponding tissues (Figure 2D). Interestingly, overexpression of *nhr-156* is sufficient to induce expression of *cyp-33C2* even in the absence of *P. brevicompactum* (Figure 2E). It is possible that the *nhr-156* multicopy DNA array generated in these strains bypasses its physiological regulation, resulting in ligand-independent activity. While this result precludes straight-forward interpretation of our rescue experiments, it provides further evidence that *nhr-156* can regulate *cyp-33C2::GFP* induction. Furthermore, *nhr-156* overexpression in AMsh glia induces *cyp-33C2::GFP* expression only in AMsh glia, demonstrating a cell-autonomous function. Consistent with their cell-autonomous functions, expression of *nhr-45* and *nhr*-*156* is observed in intestinal cells and AMsh glia, as well as in other cell types (Figure 2F,G). In these studies, *nhr-45* expression was monitored using a recombineered fosmid encoding a translational fusion to GFP, revealing NHR-45::GFP accumulation in nuclei (Figure 2F). *nhr-156* expression was observed using an *nhr-156* promoter::GFP transcriptional reporter (Figure 2G).

### *nhr-45* is a major regulator of *P. brevicompactum*-dependent XME induction

To determine if additional XME genes induced by *P. brevicompactum* are dependent on *nhr-45*, we aimed to carry out RNAseq experiments on *nhr-45* mutant animals. These studies were complicated by our observation (see below) that *P. brevicompactum* at high concentrations is toxic to *nhr-45* mutant animals and causes developmental defects. To circumvent this issue, we generated an *nhr-45* mutant line with a genomically-integrated intestine-specific *nhr-45*-expressing transgene that exhibits only minimal rescue of *cyp-33C2::GFP* induction, but is nonetheless able to rescue the developmental defects. We predicted that at least some *nhr-45* dependent transcripts could be identified in this strain, although the extent of this dependence may be underestimated.

By comparing *nhr-45* mutant animals to wild-type animals following exposure to *P. brevicompactum*, we observed that 31 of the 100 *P. brevicompactum*-induced genes we identified require *nhr-45* for their full induction (fold-decrease >2, p-adj < 0.05 when comparing *nhr-45* mutant to wild-type animals) (Table S1). Of the 18 inducible XMEs we identified, 14 are *nhr-45*-dependent (Figure 2H), including *cyp-33C2*, as predicted. We validated the RNAseq results by assessing induction of *cyp-14A4p::GFP* in the intestine and in AMsh glia, and found that in both tissues induction was indeed *nhr-45*-dependent (Figure S1B). With a few exceptions, basal expression of XMEs in the absence of *P. brevicompactum* was not significantly affected in *nhr-45* mutants (Figure 2H). These results show that *nhr-45* is a major regulator of *P. brevicompactum*-dependent XME induction in both the intestine and AMsh glia.

### *P. brevicompactum* is toxic for *nhr-45* mutant animals that fail to induce XMEs in the intestine

Our transcriptome analysis of *P. brevicompactum-*inducible genes revealed strong enrichment for stress response genes involved in detoxification. Furthermore, a large number of mycotoxins have been shown to have nematocidal activities (Li et al., 2007). Together, these observations suggest that *P. brevicompactum* may have toxic effects on *C. elegans*, and that induction of detoxification enzymes may represent a stress response aimed at clearing mycotoxins. Supporting this idea, we observed that *nhr-45* mutants grown on a plate densely seeded with *P. brevicompactum* develop slowly, and in some cases arrest and die at early larval stages. This is in contrast to wild-type animals, which are able to tolerate *P. brevicompactum* exposure (Figure 3A,B). Importantly, *nhr-45* mutant animals show normal developmental profiles in the absence of mold (Figure 3A), indicating that *nhr-45* does not generally affect development. The developmental defect resulting from *P. brevicompactum* exposure can be rescued by restoring *nhr-45* expression in the intestine, but not in AMsh glia (Figure 3C).

**Figure 3.**
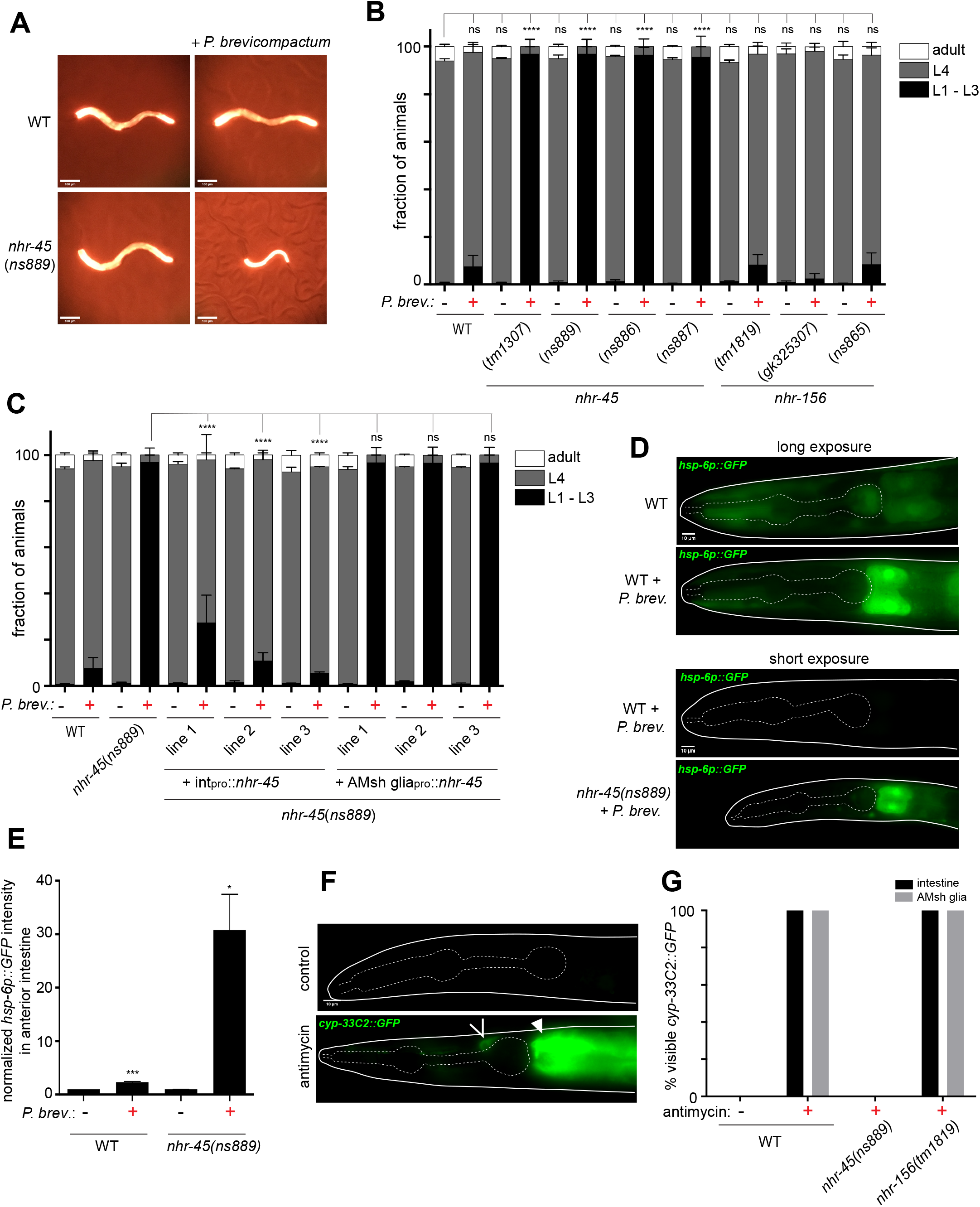
*P. brevicompactum* exposure causes mitochondrial stress and is toxic for *nhr-45* mutants with impaired XME induction. (A) Representative images of indicated strains grown for 2 days from L1 arrest with or without *P. brevicompactum* exposure. *elt-2p::mCherry* is used as a marker to visualize animals. (B,C) Scoring of developmental stages of indicated strains grown as in (A). (B) **** p < 0.0001 (L4 stage, ANOVA vs WT without *P.brev.*). (C) **** p < 0.0001 (L4 stage, ANOVA vs *nhr-45(ns889)*+*P. brev.*). (D) *hsp-6p::GFP* expression under the indicated conditions. (E) Quantification of *hsp-6p::GFP* fluorescence intensity in anterior intestine, normalized to WT without *P. brevicompactum* exposure. *** p < 0.001, * p < 0.05 (t-test vs corresponding line without *P. brev.*). (F) expression of *cyp-33C2::GFP* following antimycin treatment. Open arrowhead indicates AMsh glia, closed arrowhead indicates intestine. (G) Scoring of *cyp-33C2::GFP* expression in intestine and AMsh glia following antimycin treatment.

*nhr-45* was previously shown to regulate XME induction following mitochondrial stress during *Pseudomonas aeruginosa* infection (Mao et al., 2019). To assess whether mitochondrial stress plays a role in *P. brevicompactum* toxicity, we measured *hsp-6p::GFP* expression (Melber and Haynes, 2018). Wild-type animals exposed to *P. brevicompactum* show a modest but significant increase in *hsp-6* expression in the anterior cells of the intestine, compared to animals cultured without *P. brevicompactum* (Figure 3D,E). *nhr-45* mutant animals, which fail to induce XME expression and show signs of toxicity on *P. brevicompactum*, show massive elevation of *hsp-6::GFP* in the anterior gut cells (Figure 3D,E). These results suggest that mycotoxins produced by *P. brevicompactum* cause mitochondrial stress, which can be mitigated by inducing expression of *nhr-45*-dependent detoxifying enzymes. Supporting this idea, we found that applying a known mitochondrial inhibitor, antimycin, phenocopies exposure to *P. brevicompactum*, resulting in *cyp-33C2::GFP* induction in the intestine in an *nhr-45*-dependent manner (Figure 3F,G). Furthermore, we observed that *nhr-45* mutants are hypersensitive to the toxic effects of antimycin, consistent with previous observations (Mao et al., 2019). Interestingly, antimycin also induced *cyp-33C2::GFP* in AMsh glia, and this also requires *nhr-45*. However, in contrast to the effect of *P. brevicompactum*, antimycin-dependent induction of *cyp-33C2* in AMsh glia does not require *nhr-156*, suggesting that additional pathways, other than mitochondrial stress pathways, may underlie the effect of *P. brevicompactum* on AMsh glia (Figure 3F,G).

Together, our results support a model in which the mold *P. brevicompactum* produces mycotoxins that target mitochondria in the exposed anterior intestinal cells of *C. elegans*, and that *nhr-45*-dependent detoxification pathways are required to clear these toxins and allow animals to tolerate moldy environments.

### Natural variation in *nhr-156* underlies phenotypic diversity in AMsh glial transcriptional responses to microbes

Induction of XMEs has been described following exposure of *C. elegans* to several model pathogens (Engelmann et al., 2011). We used the *cyp-33C2::GFP* reporter to assess whether bacterial pathogens also induce XME expression in AMsh glia. We found that the *Serratia marcescens* strain Db10 induces expression of *cyp-33C2* in AMsh glia (as well as in uncharacterized cells near the anterior bulb of the pharynx), and that this induction also requires *nhr-45* and *nhr-156* (Figure 4A,B).

**Figure 4.**
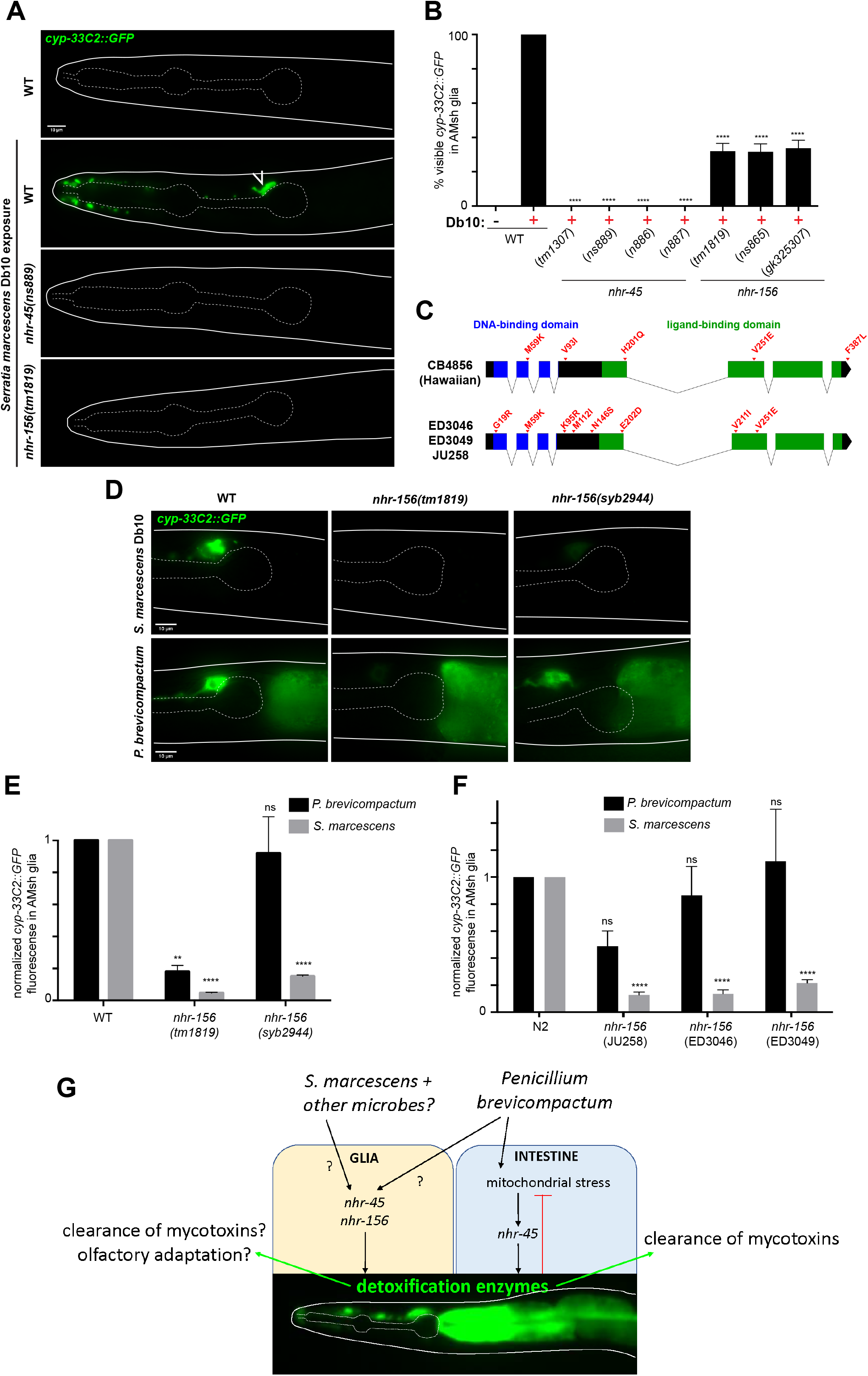
Natural variations in *nhr-156* underlie phenotypic diversity in AMsh glial transcriptional responses to microbes. (A) *cyp-33C2::GFP* expression following *S. marcescens* Db10 exposure. Open arrow indicates AMsh glia. (B) scoring of *cyp-33C2::GFP* induction in AMsh glia. **** p < 0.0001 (ANOVA vs WT + Db10) (C) schematic showing SNPs in the *nhr-156* gene that cause amino-acid changes. (D) *cyp-33C2::GFP* expression in AMsh glia following exposure to *S. marcescens* or *P. brevicompactum* in the indicated strains. *syb2944*, WT N2 background with the endogenous *nhr-156* locus replaced with the Hawaiian version of the gene. (E,F) Quantification of *cyp-33C2::GFP* fluorescence in AMsh glia, normalized to WT for each condition. Genotypes in (F) refer to representative F2 recombinant lines that were generated by crossing *cyp-33C2::GFP* from N2 into the indicated isolates. **** p < 0.0001, ** p < 0.01 (ANOVA vs corresponding WT).

The NHR family in *C. elegans* has undergone extensive expansion and diversification, and this has led to the idea that divergent NHRs may have evolved to allow adaptations to environmental conditions (Arda et al., 2010; Robinson-Rechavi et al., 2005; Sladek, 2011; Taubert et al., 2011). We noticed that there are a number of single nucleotide polymorphisms (SNPs) in *nhr-156* sequences of several distinct isolates of *C. elegans*, including the Hawaiian strain CB4856, compared to the reference N2 Bristol strain. These SNPs result in predicted coding changes in both the DNA-binding and ligand-binding domains (Figure 4C). To assess whether these changes are functionally important, we carried out a series of experiments to determine whether the Hawaiian variant of *nhr-156* exhibits alterations in its microbe-dependent activity. We initially crossed the *cyp-33C2::GFP* reporter from the N2 background into CB4856, and were surprised to readily isolate recombinant F2 lines in which *cyp-33C2::GFP* induction by *Serratia marcescens* was lower than observed in N2 (Figure S2A). Partial SNP mapping of the recovered recombinant lines revealed 100% linkage to the Hawaiian variant of *nhr-156* (16 of 16 F2 lines). This result suggests that there is a locus near the Hawaiian *nhr-156* gene that reduces *cyp-33C2::GFP* induction in AMsh glia following *Serratia marcescens* exposure. The Hawaiian locus responsible for this phenotype failed to complement *nhr-156* loss-of-function alleles, suggesting that *nhr-156* itself is the relevant gene.

To confirm these results, we generated the *nhr-156(syb2944)* strain in which the N2 *nhr-156* locus is replaced with the Hawaiian variant of the gene. Remarkably, this single-gene substitution phenocopies AMsh glia transcriptional defects observed in the Hawaiian recombinant lines (Figure 4D,E). Intriguingly, N2 animals bearing the *nhr-156(syb2944)* allele still show normal *cyp-33C2::GFP* induction in AMsh glia following exposure to *P. brevicompactum*, demonstrating that the Hawaiian variant of the gene differentially affects the response to different microbes (Figure 4D-E).

We also examined three additional *C. elegans* wild isolates with SNPs in the *nhr-156* locus (Figure 4C). Each isolate generated apparently mutant recombinant F2 lines when crossed with the *cyp-33C2::GFP* reporter and exposed to *Serratia marcescens* (Figure S2A); again with 100% linkage between the phenotype and the variant *nhr-156* locus in each case (16 of 16 F2 lines for each isolate). These recombinant lines also show relatively normal *cyp-33C2* induction following

*P. brevicompactum* exposure (Figure 4F). As a control, we obtained two wild isolates that do not harbor codon-changing SNPs in the *nhr-156* locus (JU1088 and JU1171), and found that these strains generate only fully wild-type recombinant F2 lines when crossed with the *cyp-33C2::GFP* marker. Together these results demonstrate that divergent sequences in the *nhr-156* gene found in *C. elegans* strains isolated from distinct geographical locations result in differential transcriptional responses to microbe exposure, supporting the idea that the expansion and diversification of the NHR family underlies evolutionary adaptation to environmental conditions.

## DISCUSSION

In this study, we characterized a novel interaction between *C. elegans* and *P. brevicompactum*, an ecologically-relevant species of mold that has been found in substrates that harbor wild isolates of *C. elegans* (Dirksen et al., 2016). We showed that *P. brevicompactum* causes mitochondrial stress in the intestine, accompanied by *nhr-45*-dependent induction of stress response genes. Failure to induce expression of detoxifying enzymes in the intestine correlates with toxicity to the mold, suggesting that this pathway evolved to allow *C. elegans* to tolerate toxic molds it encounters in its natural habitats (Figure 4G). This model is consistent with a recent study that demonstrated a role for *nhr-45* in the induction of XME expression following mitochondrial dysfunction (Mao et al., 2019). Our data supports a general model of pathogenesis in *C. elegans*, in which microbial toxins disrupt core cellular machineries, and this cellular damage triggers host stress responses (Liu et al., 2014; Melo and Ruvkun, 2012).

XMEs in *C. elegans* have primarily been studied in the intestine (Hartman et al., 2021). In this study we have demonstrated that XMEs are also induced in AMsh glia following *P. brevicompactum* or *Serratia marcescens* exposure. To our knowledge, this is the first demonstration of such induction in glia. While the implications of microbe-dependent XME induction in glia remain under investigation, it is notable that high levels of XME expression are observed in sustentacular glial cells in the mammalian olfactory epithelium (Heydel et al., 2013), as well as in support cells of insect antennae (Leal, 2013), where they have been proposed to regulate olfactory signaling by modifying odorant molecules. Given the essential role AMsh glial cells play in regulating chemosensory behavior (Bacaj et al., 2008; Wallace et al., 2016), it is fascinating to speculate that XME induction in AMsh glia may regulate the behavioral adaptations that accompany exposure to pathogens (Kim and Flavell, 2020). We note, however, that we did not see defects in aversive conditioning to *Serratia marcescens* in *nhr-45* or *nhr-156* mutants when assessed by lawn-leaving assays (Figure S2).

The NHR family of *C. elegans* has undergone expansion as well as sequence diversification, particularly in the ligand-binding domain (Robinson-Rechavi et al., 2005). This has led to the idea that environmental cues rather than endogenous endocrine signals may constitute ligands for many NHRs, and that divergent NHR sequences in free-living nematodes may have evolved to allow animals to adapt to environmental conditions (Sladek, 2011; Taubert et al., 2011). Our observation that natural variations in the *nhr-156* gene in wild isolates result in differential transcriptional responses in AMsh glia to microbe exposure supports this idea. Future work will investigate the molecular basis for these differences in *nhr-156* activity. It is intriguing that the variants share a common amino acid polymorphism in the ligand-binding domain, raising the possibility that this change underlies differential activation in response to microbial cues from the environment that act as ligands.

## ACKNOWLEDGEMENTS

We thank Shaham lab members for helpful discussions and the Rockefeller University Genomics Resource Center for outstanding technical support. Some strains were provided by the CGC, which is funded by NIH Office of research Infrastructure Programs (P40 OD010440). We thank the National Bioresource Project (NBRP), Cornelia Bargmann, Jonathan Ewbank and Howard Hang for strains. S.W. was a William N. and Bernice E. Bumpus Foundation postdoctoral fellow. This work was supported in part by NIH grant R35 NS105094 to S.S.

## AUTHOR CONTRIBUTIONS

S. W. – conceptualization, methodology, validation, formal analysis, investigation, data curation, writing, visualization, supervision, project administration. S. S. – conceptualization, methodology, software, formal analysis, resources, writing, supervision, project administration, funding acquisition. M. L. – conceptualization, methodology, validation, investigation, data curation. H. H. – software, formal analysis, data curation. Y. L. – methodology, software, formal analysis, resources, data curation, supervision, project administration.

---

**Figure S1. Related to Figure 2.**
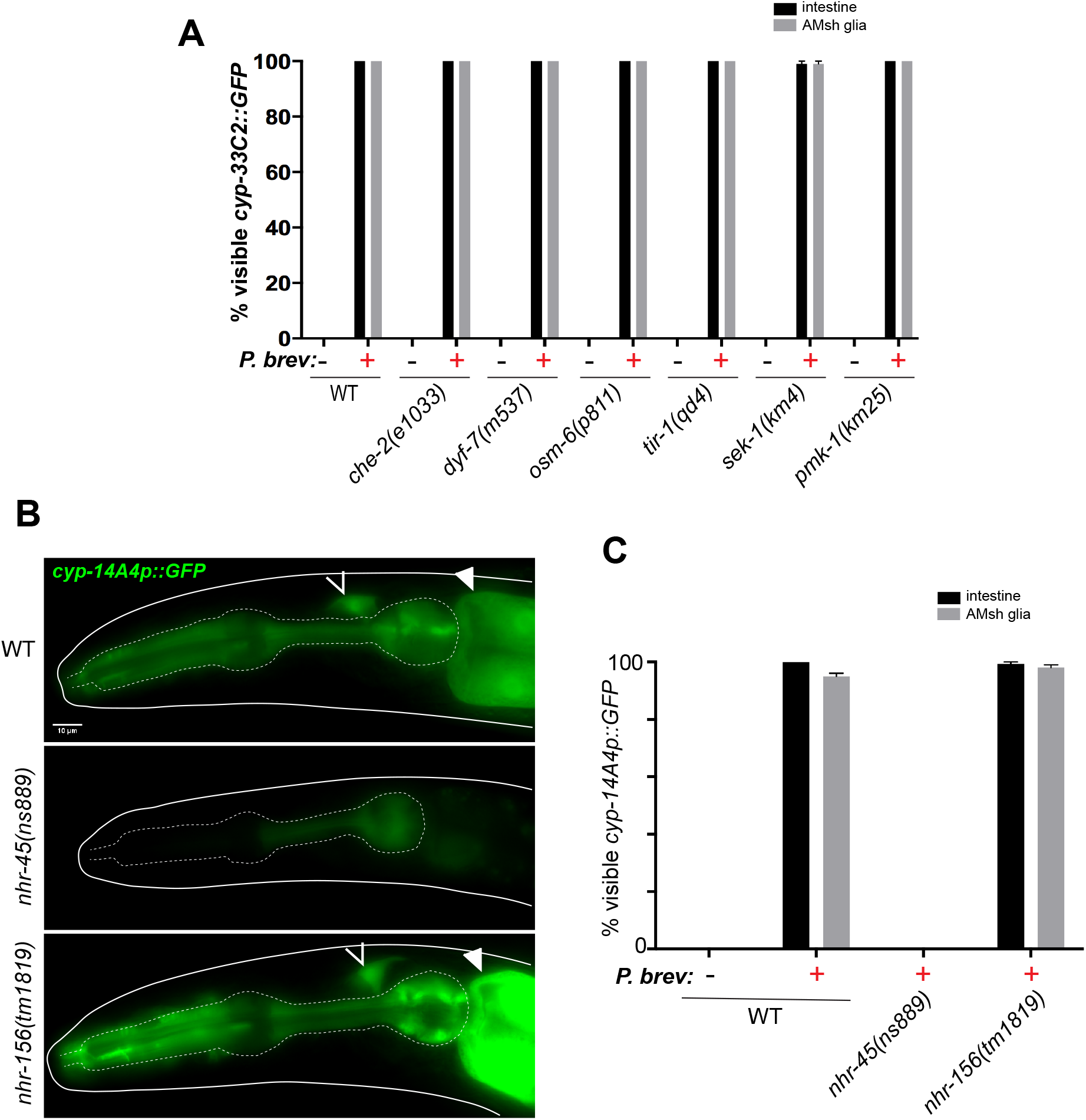
(A) *cyp-33C2::GFP* induction following *P. brevicompactum* exposure in the indicated mutant strains. (B) *cyp-14A4p::GFP* induction following *P. brevicompactum* exposure. Open arrowhead, AMsh glia. Closed arrowhead, intestine. (C) Quantification of mutant strains from (B).

**Figure S2. Related to Figure 4.**
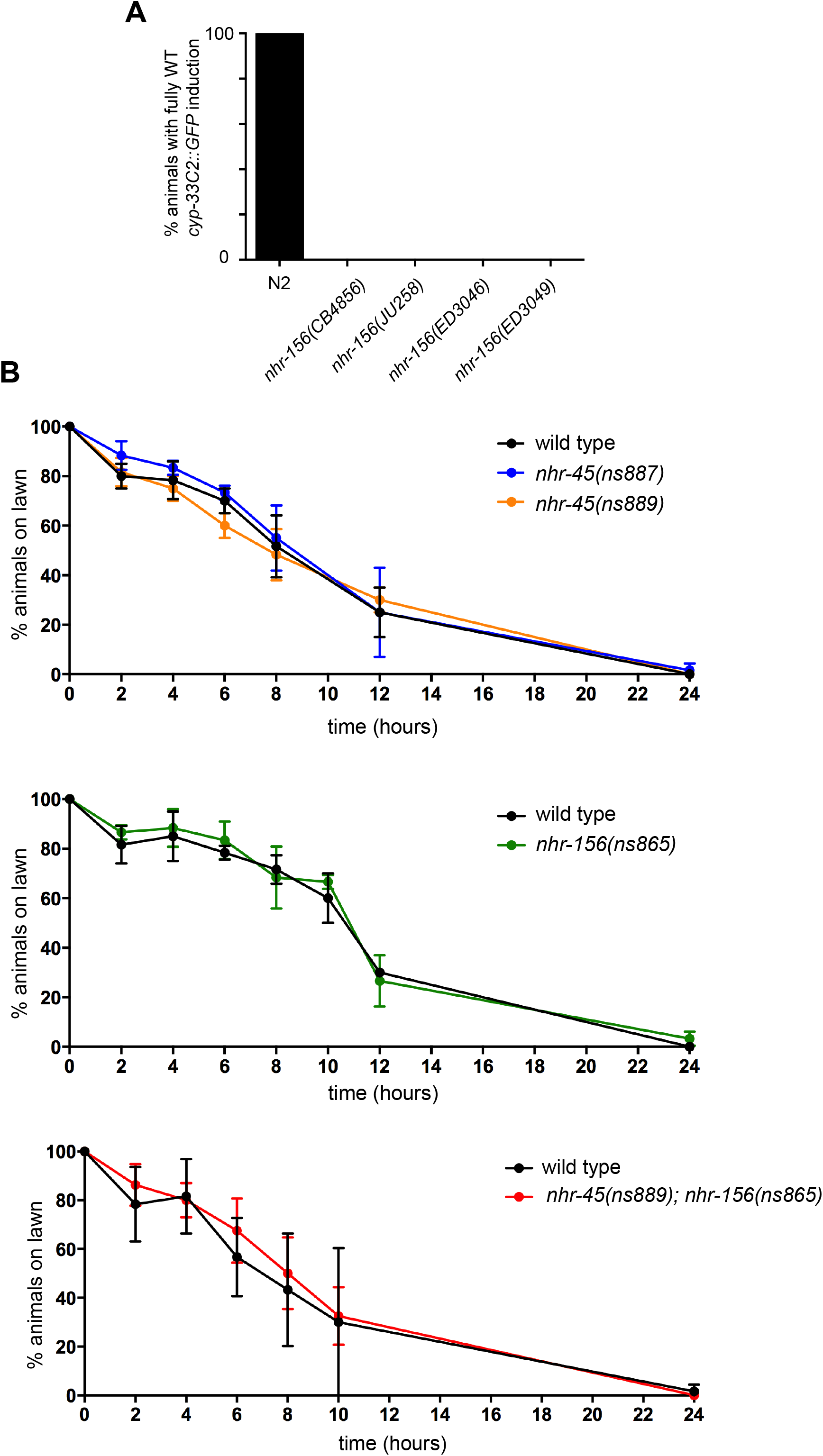
(A) *cyp-33C2::GFP* induction by *Serratia marcescens* Db10 in AMsh glia was quantified by scoring the percentage of animals that show fully wild-type levels of expression. Genotypes refer to representative F2 recombinant lines that were generated by crossing *cyp-33C2::GFP* from N2 into the indicated isolates (B) *Serratia marcescens* Db10 lawn-leaving assays for the indicated genotypes.

## METHODS

### Strains and Plasmids

*C. elegans* strains were maintained using standard methods (Stiernagle, 2006). Wild-type strain is Bristol N2.

#### Alleles used in this study

*nhr-45(tm1307); nhr-156(tm1819); nhr-156(gk325307); che-2(e1033); dyf-7(m537); osm-6(p811); tir-1(qd4); sek-1(km4); pmk-1(km25)*.

#### Alleles generated in this study

*nhr-45(ns886)* (14 bp deletion in exon1, generated by CRISPR); *nhr-45(ns887)* (11 bp insertion in exon 1, generated by CRISPR); *nhr-45(ns889)* (7 bp deletion in exon 1, generated by CRISPR); *nhr-156(ns865)* (Gln29>ochre, generated by EMS mutagenesis); *nhr-156(syb2944)*(endogenous *nhr-156* gene removed from N2 background and replaced with the Hawaiian variant of the gene – strain created by SunyBiotech).

#### Transgenes used in this study

*mgIs73* (*cyp-14A4p::GFP::cyp-14A4 3’UTR* + *myo-2p::mCherry*); *stIs11634* (*ugt-1::H1-mCherry* + *unc-119(+)*); *jrIs1* (*cyp-13A7p::GFP* + *unc-119(+)*); *nsIs143* (*F16F9.3p::dsRed*); *nsIs698* (*mir-228p::NLS-RFP* + *unc-122p::RFP*); *zcIs13* (*hsp-6::GFP*)

#### Transgenes generated in this study

*nsIs775* (pSW81 + *elt-2p::mCherry*); *nsIs910* (pSW124 + *unc-122p::GFP*)

#### Extrachromosomal arrays used in this study

*otEx1182* {g*st-44p*::*gst-44*::*GFP* + *rol-6(dom)}*

#### Extrachromosomal arrays generated in this study

*nsEx5944* {pSW74 + *elt-2p::mCherry*}; *nsEx6240* {pSW124 + *unc-122p::GFP*, line 1}; *nsEx6241* {pSW124 + *unc-122p::GFP*, line 2}; *nsEx6242* {pSW124 + *unc-122p::GFP*, line 3}; *nsEx6238* {pSW110 + *unc-122p::GFP*, line 1}; *nsEx6239* {pSW110 + *unc-122p::GFP*, line 2}; *nsEx6240* {pSW110 + *unc-122p::GFP*, line 3}; *nsEx6194*{pSW118 + *unc-122p::GFP*, line 1}; *nsEx6196*{pSW118 + *unc-122p::GFP*, line 2}; *nsEx6197*{pSW118 + *unc-122p::GFP*, line 3}; nsEx6176 {WRM0636C_B02(pRedFlp-Hgr)(nhr-45[15070]::S0001_pR6K_Amp_2xTY1ce_EGFP_FRT_rpsl_neo_FRT_3xFlagdFRT::unc-119-Nat (*nhr-45* fosmid recombineered with GFP) + *elt-2p::mCherry*}; *nsEx6180* {pSW121 + *elt-2p::mCherry*}.

#### Plasmids generated in this study

pSW81 (*cyp-33C2*p(2kb)::c*yp-33C2*::GFP::*unc-54 3’UTR*); pSW124 (*elt-2p::nhr-45 cDNA::unc-54 3’UTR*); pSW74 (*gst-33p::gst-33::GFP::unc-54 3’ UTR*); pSW121 (*nhr-156p::GFP::unc-54 3’ UTR*); pSW110 (*F16F9.3p::nhr-45 cDNA::unc-54 3’UTR*); pSW118 (*F16F9.3p::nhr-156 cDNA::unc-54 3’UTR*)

### *Penicillium brevicompactum* culture

*P. brevicompactum* (ATCC catalog number 9056) was maintained on NGM plates. Cultures were kept at room temperature for 2 weeks or at 4 C for 2 months. Fresh cultures were created by streaking spores from an existing culture. Mold species were genotyped by scraping material from a plate in M9 buffer, briefly centrifuging to pellet material, and lysing in worm lysis buffer (10 mM Tris pH 8.0, 50 mM KCl, 2.5 mM MgCl_2_, 0.45% NP-40, 0.45% Tween-20, 0.01% gelatin, 200 μg/ml proteinase K) by incubating at 65 C for 1 hour. Genotyping PCRs were performed directly on lysates using ITS, LSU and SSU primers (Schoch et al., 2012).

To expose *C. elegans* to *P. brevicompactum*, spores were scraped from a culture plate and added to NGM liquid cultures immediately prior to pouring NGM plates. Plates were left to dry for several days, before being seeded with *E. coli* OP50. Mold spores were calibrated such that approximately 50% of the surface of the plate was covered by mold when the plates were used for *C. elegans* assays. Unless otherwise stated, assays were performed by plating synchronized L1s on to moldy plates and scoring phenotypes after 2 days of growth at 20 C.

### RNA extraction and Sequencing

*C. elegans* were prepared for transcriptome analysis by culturing on plates with or without *P. brevicompactum*, as described above. To avoid obtaining artifacts due to differences in developmental staging, it is critical that are samples are staged correctly. RNA was extracted from samples approximately 48 hours from L1 arrest, at which time point the majority of animals had reached the young adult stage. The precise timing was decided empirically for each condition in each experiment, such that some plates were left for up to 52 hours if they contained an appreciable number of animals that were still at the L4 stage at the 48 hour time point. To overcome the toxicity of *P. brevicompactum* for *nhr-45* mutant animals, we took advantage of the transgenic line *nsIs910* (*elt-2p::nhr-45 cDNA*) that contains a partial rescue of *nhr-45* function in the intestine. This line showed only minimal rescue of *cyp-33C2* induction in the intestine, but showed relatively normal developmental timing on *P. brevicompactum.* We hypothesized that *nhr-45* induces multiple redundantly-acting XME targets, and that the partial rescue of *nhr-45* function was not sufficient to rescue *cyp-33C2* induction, but was sufficient to prevent toxicity due to its regulation of other targets. We therefore hypothesized that this line would be a useful tool to identify *nhr-45* target genes, albeit with the caveat that we would likely underestimate the extent to which XME expression depends on *nhr-45*. The sequencing data we obtained confirmed this hypothesis. Many of the inducible XMEs required *nhr-45* for their full induction, but some still showed partial induction due to the partial rescue. This line was therefore useful for identifying *nhr-45* target genes without encountering developmental staging problems.

To extract total RNA, animal pellets were suspended in 4x volume Trizol-LS, vortexed for 5 minutes and subjected to two rounds of freeze-thawing with additional vortexing after each thaw. RNA was then isoted using chloroform extraction following the manufacturer’s guidelines. Total RNA was then subjected to RNeasy column cleanup with DNAse treatment (company). cDNA library preparation and sequencing were as performed by The Rockefeller University Genomics Research Center, using Illumina TruSeq stranded mRNA-Seq library preparation kit with polyA selection, and Illumina NextSeq500.

### RNA-Seq Quality Assessment and Differential Gene Expression Analysis

Sequencing data was assessed for quality control using FastQC v0.11.12 (https://www.bioinformatics.babraham.ac.uk/projects/fastqc/) with default parameters. Trimmed fastq files were aligned to WBcel235 using STAR v2.4a aligner with default parameters(Dobin et al., 2013). The alignment results were then evaluated through Qualimap v2.2(García-Alcalde et al., 2012) to ensure that all the samples had a consistent coverage, alignment rate, and no obvious 5’ or 3’ bias. Aligned reads were then summarized through featureCounts on gene (Ensemble gene model Caenorhabditis_elegans.WBcel235.103.gtf). To eliminate the effect of library size, summarized count matrix was normalized through edgeR v 3.28.1 (McCarthy et al., 2012; Robinson et al., 2010). Voom from limma v 3.42.2 was applied to estimate the fold change (Ritchie et al., 2015). A BH-adjusted *p*-value of less than 0.05 (p-adj < 0.05) was used to select genes that have a significant expression change.

### RNA Interference

RNAi by feeding was carried out as described previously (Kamath and Ahringer, 2003), with modifications to allow exposure to *P. brevicompactum*. Overnight cultures of *E. coli* HT115 carrying pL4440 vectors targeting specific genes (Ahringer and Vidal libraries) were seeded on to *P.brevicompactum*-NGM plates (described above) supplemented with carbenicillin (25 ug/ml) and IPTG (1 mM). *C. elegans* were added to plates as L1 larvae, and scored 48 hours later.

### Fluorescence Microscopy

Imaging was performed using an Axioplan II fluorescence microscope equipped with an AxioCam camera. Images were processed using Image J software. Processing involved orienting the samples, cropping to show the region of interest, and setting the upper and lower intensity thresholds for display. All images in a given experiment were processed in an identical manner. For fluorescence quantification, a region of interest was defined and mean pixel intensity was measured using Image J software. Background subtractions was performed by measuring the mean pixel intensity of a corresponding background region. At least 20 animals were imaged for each sample in a given replicate, and at least 3 independent replicates were imaged for each experiment. Data is presented for each condition as the mean of the independent replicates normalized to the control for each replicate.

### *Serratia marcescens* lawn-leaving assays

20 young adults were picked on to freshly seeded lawns of *Serratia marcescens* Db10 per replicate, and the percentage of animals occupying the lawn was calculated at each time point.

